# Enhanced Tactile Coding in Rat Neocortex Under Darkness

**DOI:** 10.1101/2025.04.11.648316

**Authors:** Kotaro Yamashiro, Shiyori Tanaka, Nobuyoshi Matsumoto, Yuji Ikegaya

## Abstract

Sensory systems are known for their adaptability, responding dynamically to changes in environmental conditions. A key example of this adaptability is the enhancement of tactile perception in the absence of visual input. Despite behavioral studies showing visual deprivation can improve tactile discrimination, the underlying neural mechanisms, particularly how tactile neural representations are reorganized during visual deprivation, remain unclear. In this study, we explore how the absence of visual input alters tactile neural encoding in the rat somatosensory cortex (S1). Rats were trained on a custom-designed treadmill with distinct tactile textures (rough and smooth), and local field potentials (LFPs) were recorded from S1 under light and dark conditions. Machine learning techniques, specifically a convolutional neural network, were used to decode the high-dimensional LFP signals. We found that the neural representations of tactile stimuli became more distinct in the dark, indicating a reorganization of sensory processing in S1 when visual input was removed. Notably, conventional amplitude-based analyses failed to capture these changes, highlighting the power of deep learning in uncovering subtle neural patterns. These findings offer new insights into how the brain rapidly adapts tactile processing in response to the loss of visual input, with implications for multisensory integration.

## Introduction

The primary somatosensory cortex (S1) is integral to the encoding of tactile information, processing sensory inputs from various regions of the body (Delhaye et al., 2018; Di Plinio et al., 2020; Piras et al., 2020; Serino, 2019). This cortical area is crucial for our sense of touch, facilitating the perception and interpretation of sensations such as pressure, vibration, temperature, and pain (Bushnell et al., 1999; Luna et al., 2005; Moulton et al., 2012). These external stimuli are represented by dynamic neural activity in S1, enabling animals to discriminate distinct sensory experiences (Bensmaia et al., 2008; Goodwin and Wheat, 2004; Koch and Fuster, 1989; Salinas et al., 2000). Crucially, neural representations in S1 are highly adaptable, shaped not only by feedforward mechanisms but also by attention, motivation, and inputs from other sensory modalities (Butler et al., 2012; Driver and Spence, 2000; Eimer and Forster, 2003; Schürmann et al., 2004; Ziegler et al., 2023). This adaptability facilitates rapid adjustments to changing environmental contexts, supporting survival and optimal sensory processing (Abraira and Ginty, 2013; Dijkerman and de Haan, 2007).

One prominent example of such adaptability is cross-modal processing, wherein the absence or reduction of visual input enhances tactile abilities. (Bulusu and Lazar, 2024; Hopkins et al., 2017; Nikbakht et al., 2018; Sugiyama et al., 2019). For instance, prolonged visual deprivation, such as blindness from an early age, results in heightened tactile discrimination abilities (Goldreich and Kanics, 2003; Norman and Bartholomew, 2011; Van Boven et al., 2000; Wong et al., 2011), driven by cortical reorganization where visual cortical areas are repurposed to support tactile processing (Burton, 2003; Karlen et al., 2006; Sadato et al., 1996). Notably, however, tactile performance improvements are also observed in sighted individuals during short-term visual deprivation (Facchini and Aglioti, 2003; Kauffman et al., 2002; Pascual-Leone and Hamilton, 2001a). Such individuals demonstrate enhanced tactile discrimination during temporary darkness or blindfolding, indicating that rapid, context-dependent compensatory mechanisms occur even without long-term structural changes (Bola et al., 2017; Boroojerdi et al., 2000; Merabet et al., 2008). Thus, although tactile enhancement has been studied extensively in the context of long-term sensory deprivation, the neural mechanisms underlying short-term, context-dependent modulation remain unclear.

The neural representation of tactile stimuli within S1 involves complex, distributed activity patterns, posing significant challenges for conventional analyses that primarily measure signal amplitude. Thus, the subtle, high-dimensional neural dynamics that may underlie immediate adaptations to temporary visual deprivation are difficult to detect using traditional methods. Addressing this gap requires refined experimental paradigms and advanced analytical tools to capture how tactile representations reorganize on short timescales.

To examine short-term reorganization in tactile processing, we used a behavioral paradigm using rats trained to walk naturally on a treadmill featuring distinct tactile textures (Yamashiro et al., 2024). . By manipulating visual input (light *vs*. dark conditions) while recording local field potentials (LFPs) from S1, we aimed to characterize rapid shifts in tactile neural representations. LFPs provide a rich, high-dimensional signal reflecting both synchronous and asynchronous activity across cortical populations. Concurrently, we applied deep learning techniques to decode subtle differences in neural activity patterns that traditional amplitude-based analyses could not detect. We hypothesized that temporary visual deprivation would enhance the neural distinction between tactile stimuli, leading to more differentiated representations of textures in the dark condition.

By investigating these rapid, context-dependent shifts in neural coding, this study provides valuable insights into the brain’s capacity for sensory adaptation. Our findings not only shed light on how visual deprivation influences tactile processing in the short term, but also emphasize the potential of deep learning techniques in uncovering nuanced neural reorganization.

## Materials and Methods

### Animal ethics

Animal experiments were performed with the approval of the Animal Experiment Ethics Committee at the University of Tokyo (approval numbers: P29–7 and P4–15) and according to the University of Tokyo guidelines for the care and use of laboratory animals. These experimental protocols were carried out following the Fundamental Guidelines for the Proper Conduct of Animal Experiments and Related Activities of the Academic Research Institutions (Ministry of Education, Culture, Sports, Science and Technology, Notice No. 71 of 2006), the Standards for Breeding and Housing of and Pain Alleviation for Experimental Animals (Ministry of the Environment, Notice No. 88 of 2006) and the Guidelines on the Method of Animal Disposal (Prime Minister’s Office, Notice No. 40 of 1995). While our experimental protocols have a mandate to humanely euthanize animals if they exhibit any signs of pain, prominent lethargy, and discomfort, such symptoms were not observed in any of the rats tested in this study. All efforts were made to minimize the animals’ suffering.

### Behavioral paradigm

To record LFPs during natural locomotion, we employed a custom-designed, disk-shaped treadmill with a diameter of 90 cm (Figure 1A). The treadmill’s running surface was divided into two halves, each featuring a distinct texture: one side was coated with coarse sandpaper (grain #80) and the other with fine sandpaper (grain #1000). In this setup, the rat was placed on the treadmill and secured with a fabric vest (Figure 1B).

**Figure 1.**
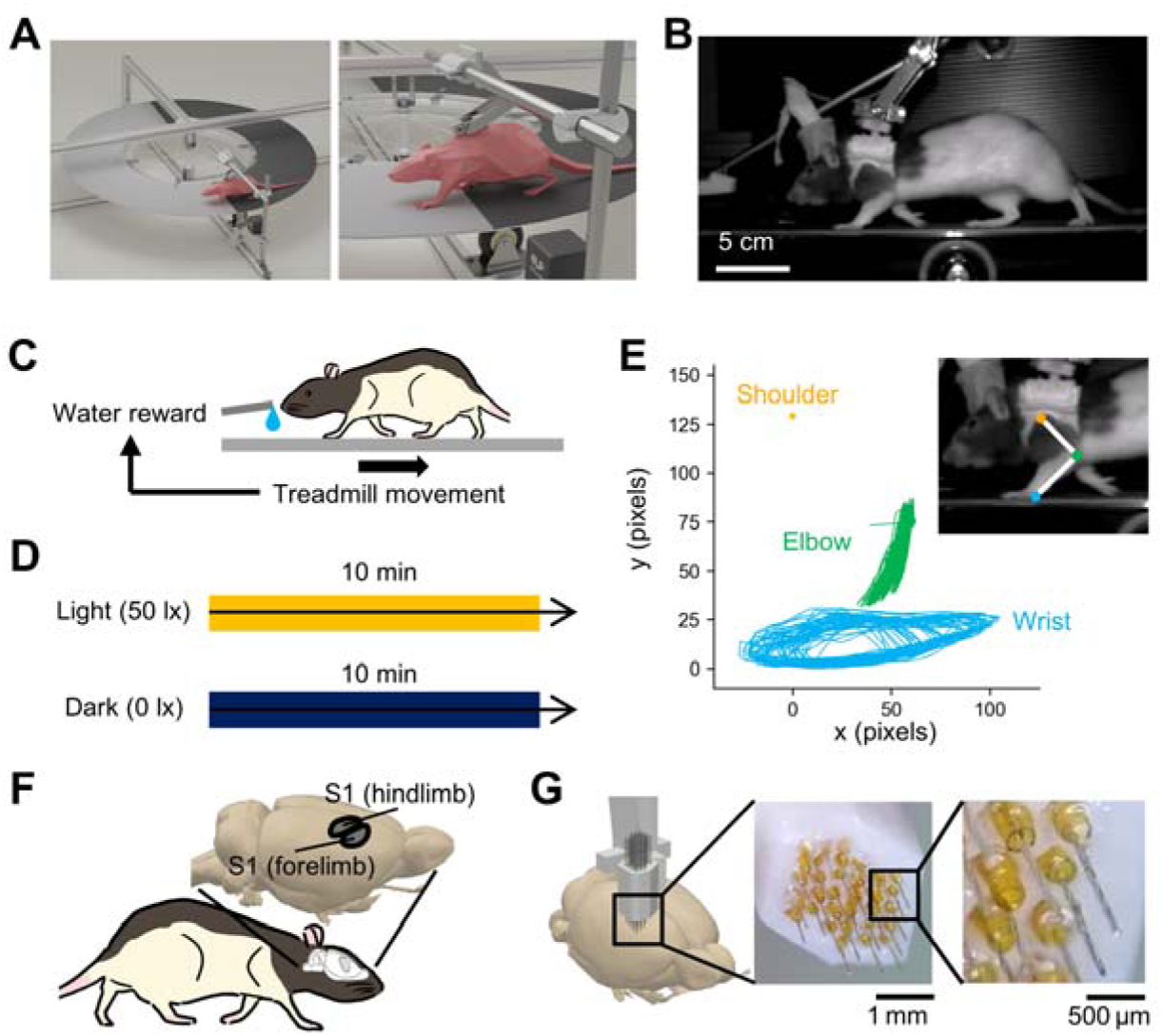
Behavioral paradigm and limb movement assessment with concurrent LFP recordings. (**A**) A diagram of the disk-shaped treadmill used in the experiment. One half of the disk is covered with #80 sandpaper, and the other half with #2000 sandpaper. (**B**) A frame from the video capturing a walking rat from a left-side perspective. (**C**) The motivation scheme. The rats were water-deprived prior to the experiments. A water port was coupled with the movement of the treadmill so that when the rat walked on the treadmill, the water would come out. This way, the rats were always motivated to walk during the whole session. (**D**) The experimental protocol, where each rat walked for 10 minutes in light (50 lx) and then for 10 minutes in darkness (0 lx). (**E**) An example trajectory of the elbow and wrist joints from one session, plotted with the shoulder joint fixed in the coordinate space (shoulder: *yellow*, elbow: *green*, wrist: *cyan*). (**F**) A schematic illustrating the forelimb and hindlimb subregions of S1. (**G**) A custom 32-channel electrode array used to record LFPs from these subregions.

Before the experiment, rats were water-restricted to ensure motivation. A waterspout was positioned in front of each rat, and its output was synchronized with the treadmill’s movement (Figure 1C). Specifically, once the treadmill began to move, 30 µL of water was dispensed from the spout, encouraging the rat to continue walking to receive its water reward.

Each rat was deprived of water until its body weight reached 85% of its baseline weight, then it was trained to walk on the treadmill for four days (one hour per day), with 30 minutes in a light environment (50 lx) followed by 30 minutes in darkness (0 lx). After successful training, the rats were allowed free access to food and water to regain their original body weight before electrode implantation surgery. Following surgical recovery, the rats were water-restricted again and placed on the treadmill to assess cross-modal interactions between the visual and tactile systems. In a single session, LFP recordings were obtained in the light environment trial, followed immediately by the dark environment trial (Figure 1D). For subset of sessions, the order of the light and dark environment was swapped, dark environment trial preceding the dark environment trial. Each trial lasted for approximately 10 minutes. Recording was performed once a day for 4-14 days.

During the trials, the trajectory of the forelimb and onsets when the forelimb came in contact with the floor were automatically identified using a previously developed deep-learning–based method (Figure 1E) (Yamashiro et al., 2024).

### Animal preparation and surgical procedures

LFPs were recorded from eleven 9- to 10-week-old Long-Evans rats (Japan SLC, Shizuoka, Japan) using a custom-designed, 32-channel electrode assembly. This assembly, fabricated from nichrome wires (761500, A-M Systems, WA, USA), targeted the right somatosensory cortex (S1) regions corresponding to the forelimb and hindlimb representations (Figure 1F). Specifically, 18 and 14 electrodes were placed in the forelimb and the hindlimb subregion respectively (Figure 1G). Each electrode tip was platinum-coated to reduce impedance to below 200 kΩ using a nanoZ tester (Plexon, TX, USA).

At the start of the surgical procedure, each rat was anesthetized with 2–3% isoflurane gas. A square craniotomy (2–6 mm posterior and 1–5 mm lateral to bregma) was then created using a dental drill. The electrode assembly was gently lowered through the cranial window to a depth of approximately 1.5 mm beneath the dura, targeting layer IV of S1. Additionally, two stainless steel screws were implanted in the bone above the cerebellum to serve as ground and reference electrodes. The recording device and electrodes were secured to the skull using stainless steel screws and dental cement. Following the surgery, each rat was housed individually in a transparent Plexiglas cage with ad libitum access to food and water for one week to ensure proper recovery.

### LFP recordings from S1

LFPs were referenced to ground, digitized at 30 kHz using the OpenEphys recording system (http://open-ephys.org) and an RHD 32-channel headstage (C3314, Intan Technologies, CA, USA), then resampled to 10 kHz for subsequent analyses (Yamashiro et al., 2020). In parallel, video was acquired at 60 Hz using a USB camera module (MCM-303NIR, Gazo, Niigata, Japan), capturing a lateral view of the rat. Each video frame was synchronized with the neural recordings using strobe signals. For a subset of recordings (n = 2 rats), speed sensors were attached to the running disk to monitor locomotor speed, which was simultaneously recorded through the OpenEphys analog input.

### Data analysis

Data was analyzed offline using custom-made scripts in Python3. For box plots, the centerline shows the median, the box limits show the upper and lower quartiles, and the whiskers cover the 10−90% percentiles. *P*L<L0.05 was considered statistically significant. All statistical *t*-tests were two-sided, and the bootstrap method was applied for multiple comparisons.

### LFP analysis

To see if there were any differences in LFPs, the LFPs were aligned to the foot strike onsets detected using deep-learning assisted methods. The aligned LFP were then categorized by the trial (light *vs.* dark) and the floor texture (smooth *vs.* rough). To analyze the amplitude of the event-related response from the foot strike, a mean trace of the aligned LFP was calculated for each channel from 32 electrodes for each condition. Foot-strike detection and all LFP alignment were intentionally forelimb-locked, because the behavioral paradigm was designed for forepaw texture contact (Figure 1E) and our array oversampled S1-forelimb (18 channels) relative to S1-hindlimb (14 channels; Figure 1G).

### A deep neural network for joint decoding of trial conditions and floor texture

A custom deep neural network (DNN) model was implemented to predict both the trial condition (*e.g.*, light *vs.* dark) and floor texture (*e.g.*, smooth *vs*. rough) from time-aligned, one-dimensional LFP segments, using the PyTorch framework. Our model architecture was inspired by one-dimensional ResNet-like structures and incorporated multiheaded outputs for simultaneous prediction of two distinct variables (He et al., 2015).

The input to the model consisted of one-dimensional LFP signals, arranged as a tensor with multiple channels (*e.g.*, 32 input channels) over time. The network began by splitting the input into two parallel convolutional pathways. The first pathway (“left” branch) applied sequential convolutional and pooling operations with relatively smaller kernel sizes and strides to incrementally reduce the dimensionality of the signal and extract fine-grained temporal features. Specifically, the model employed a two-stage convolutional process that first passed the input through a one-dimensional (1D) convolution layer with a kernel size of 7, stride of 2, and batch normalization, followed by a max pooling and an additional convolution layer. Both convolutional layers in the left branch used ReLU nonlinearities to facilitate stable and efficient feature extraction.

In contrast, the second pathway (“right” branch) processed the input through a single 1D convolutional layer with a larger kernel size (*e.g.*, 41) and a more aggressive stride (*e.g.*, stride of 8). This pathway captured broader temporal contexts from the input signals. Similar to the left branch, the right branch output was batch-normalized and passed through a ReLU activation function. After these two parallel extractions, the outputs of the left and right branches were concatenated along the channel dimension, forming a combined feature representation that integrated both fine- and coarse-grained temporal information.

The concatenated output was then passed through a max pooling operation, followed by two residual layers that employed 1D convolutional blocks (ResidualBlock) to refine feature representations. These residual layers allowed the network to learn more complex feature hierarchies by facilitating the flow of gradients during training and improving convergence, while also maintaining temporal resolution appropriate for downstream decoding.

Subsequently, the processed features were passed through an average pooling layer to summarize temporal information into a low-dimensional feature vector. This vector was flattened into a one-dimensional representation and then fed into two separate fully connected “heads” (fc_head1 and fc_head2). Each head was a simple linear layer that provided a scalar output value. By concatenating these outputs, the final layer jointly predicted two target variables from the same underlying features.

In summary, our model combined parallel convolutional branches for initial feature extraction, residual layers for robust representation learning, and multiheaded outputs to facilitate joint prediction of trial conditions and floor textures. The model was implemented in Python using PyTorch, and all parameters were optimized via standard stochastic gradient–based methods. This architecture allowed efficient and robust decoding of environmental conditions from LFP signals in both time and frequency domains.

### Training and evaluating the deep neural network

Of the 11 recorded rats, data from the initial two rats were utilized to determine the optimal model architecture. Once the architecture was established, data from the remaining 9 rats were used for training and evaluation. The raw LFP signals were resampled from 30 kHz to 10 kHz and segmented into 800 ms windows centered on the footstrike onset (*e.g.*, 400 ms before and 400 ms after). Each LFP segment was assigned two labels: one for the trial condition (*e.g.*, light *vs*. dark) and one for the floor texture (*e.g.*, smooth *vs*. rough).

For each rat, the dataset was shuffled, normalized, and then subjected to 5-fold cross-validation. In this procedure, the data were partitioned into five equal subsets; in each fold, four subsets (80%) were used for training, and the remaining subset (20%) was used for evaluation. Model training was performed for 80 epochs using a batch size of 128 and a learning rate of 10^-6^. Binary cross-entropy with logits loss (BCEWithLogitsLoss) served as the objective function. These parameters were selected to ensure stable convergence and to reduce overfitting. After the training, the model’s performance was evaluated using the evaluation dataset. The confusion matrix was calculated as the number of true positives, false positives, false negatives, and true negatives, aggregated across all predictions in the evaluation set.

### Evaluation of cluster separability between light and dark conditions

To examine how texture representations differed between light and dark conditions, we assessed cluster separability of the intermediate features extracted from the deep-learning model. Specifically, after the model was trained, we extracted the output from the penultimate layer (a 912-dimensional feature vector for each trial). Cluster separability was quantified using the silhouette score, which measures the relative distances between within-cluster and between-cluster samples.

Silhouette scores were calculated for the two texture classes (smooth and rough) within each lighting condition, using the Euclidean distance metric applied to the intermediate feature representations. For each trial, the silhouette score *s* is defined as:

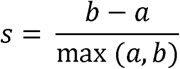

where *a* is the average distance between a point and all other points in the same cluster (intra-cluster distance), and *b* is the minimum average distance between a point and all points in the nearest different cluster (inter-cluster distance). The silhouette score, therefore, provides a normalized index of cluster separability, with larger values indicating more distinct clustering. To obtain an animal-level measure, silhouette scores were averaged across cross-validation within each rat.

### Explainability analysis using occlusion and integrated gradients

To elucidate the internal decision-making processes of the trained deep neural network (DNN), we employed two established explainability techniques: occlusion and integrated gradients.

#### Occlusion

Occlusion analysis involves systematically masking specific input features to determine their relative contribution to the model’s output (Zeiler and Fergus, 2013). In the present study, one channel was selectively occluded at a time from the 32-channel LFP input and the sensitivity of each channel was calculated. Sensitivity was defined as the corresponding change in model performance when the specified channel was occluded. By conducting these analyses for each of the nine rats, channel-specific importance scores were obtained and subsequently normalized (z-scored) to facilitate cross-subject comparisons. Channels whose removal yielded a more pronounced decrease in model performance were considered more critical for accurate prediction.

#### Integrated gradients

Integrated gradients is an attribution method that quantifies the importance of each input feature by integrating the gradient of the model’s output with respect to the input, transitioning from a baseline input to the actual input (Sundararajan et al., 2017). This approach produces class activation maps, enabling the visualization of features most influential for the model’s output. Here, integrated gradients were applied to the LFP segments for each rat, and the resulting class activation maps were averaged across subjects and trial conditions. These maps allowed us to identify salient input regions associated with both trial conditions and floor texture.

## Results

### Stable locomotion across light and dark conditions

To investigate how visual input influences tactile processing in S1, we devised an experimental paradigm where both tactile and visual inputs were independently manipulated. Rats were placed on a disk-shaped treadmill with two distinct sandpaper textures, and LFPs were recorded from walking rats. Each rat walked for 10 min in a light environment (50 lx) and then for 10 minutes in total darkness (0 lx).

To make sure that rat’s trajectory was stable across different floor textures and environmental conditions, gait parameters were extracted from the trajectories using deep-learning–based analysis (Figure 2A). From the trajectories, swing duration, stance duration, stride length, and footstrike speed were extracted. All parameters were calculated for each floor textures and environmental conditions. Comparison of all conditions revealed that none of these metrics differed significantly between floor textures or environmental conditions, indicating that overall locomotion remained stable (Figure 2B-E, *P* > 0.05, one-way analysis of variance (ANOVA) followed by TukeyLKramer *post hoc* test, *n* = 149, 149, 107 and 107 trials for smooth-light, rough-light, smooth-dark, and rough-dark, respectively). Locomotor-speed traces recorded from subset of rats. The result indicated that walking velocity was stable across trials, not affected by the lighting conditions (Supplementary Figure 1). Thus, any differences in neural activity under these conditions are unlikely to be driven by altered motor behavior.

**Figure 2.**
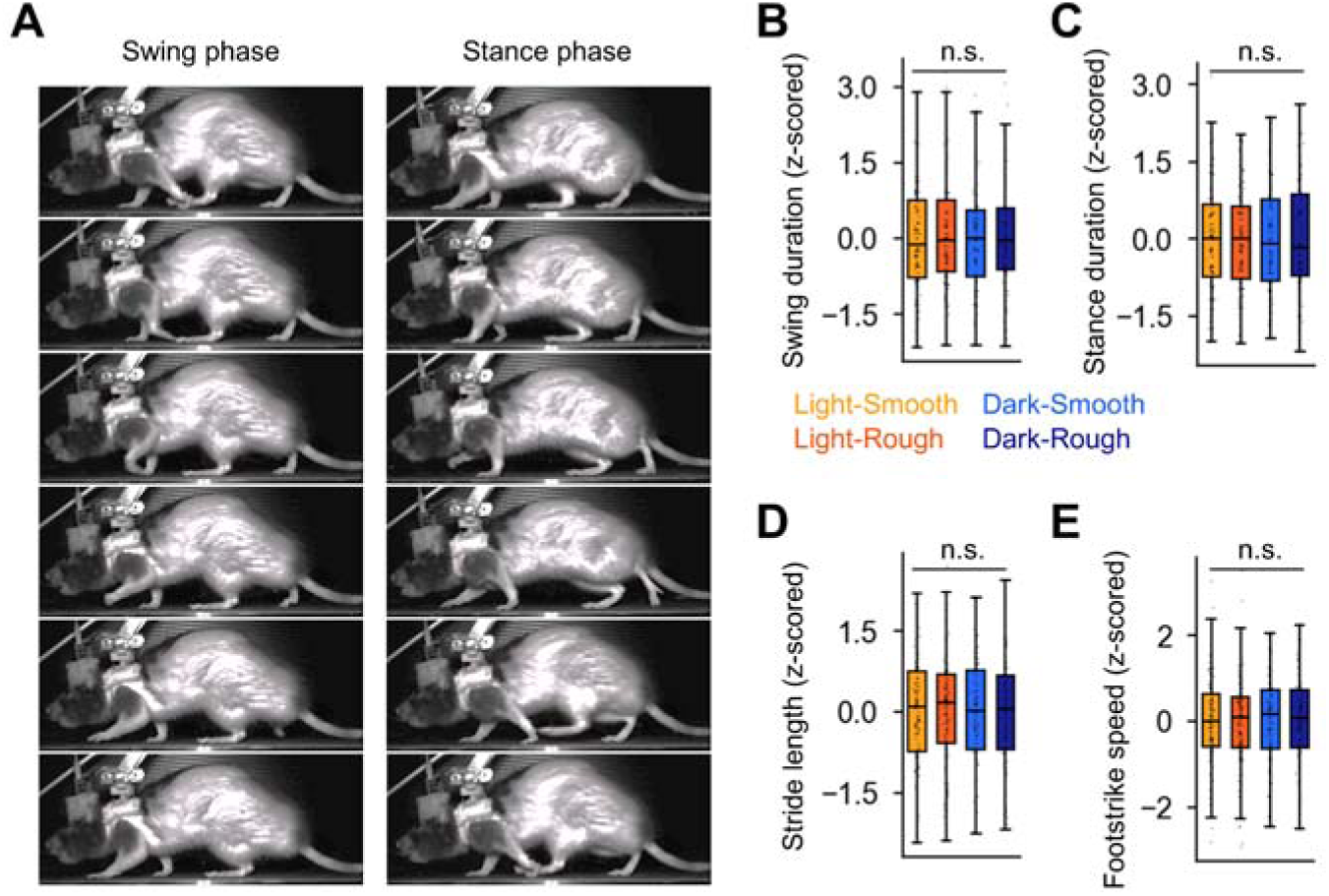
Comparison of gait parameters across textures and environmental conditions. (**A**) Swing phase *vs*. stance phase, illustrated with video frames (*left*: swing, *right*: stance). (**B**) Normalized swing duration measured for each rat under different textures (smooth *vs*. rough) and environmental condition (light *vs*. dark). Light orange and dark orange correspond to the light conditions (smooth, rough), while light blue and dark blue correspond to the dark conditions (smooth, rough). There were no significant differences among trial conditions. (**C–E**) Stance duration, stride length, and footstrike speed, respectively, under the same conditions as in B. None of these parameters differed significantly across texture types or lighting conditions.

### Characteristic of S1 LFP upon forelimb contact

We next analyzed LFPs from S1 using a custom 32-channel electrode array targeting the forelimb and hindlimb subregions (Figure 1F, G). LFP traces were first aligned to forelimb contacts with the disk surface. On inspection of a single LFP trace, no apparent event-related response could be observed (Figure 3A). However, aligning each LFP at forelimb contact with the floor, the average trace showed a clear response after the onset (Figure 3B). Across conditions, we observed a clear negative deflection in LFPs that was most pronounced in the forelimb subregion (Figure 3C). The amplitude of the response was significantly larger in all rats, consistent with the topographic specificity of S1 (Figure 3D, *P* = 1.89×10^-3^, *t*_10_ = -2.2, paired *t*-test, *n* = 11 rats). This was expected since the epochs were forelimb-locked, making larger deflections and greater importance of forelimb channels consistent with somatotopy.

**Figure 3.**
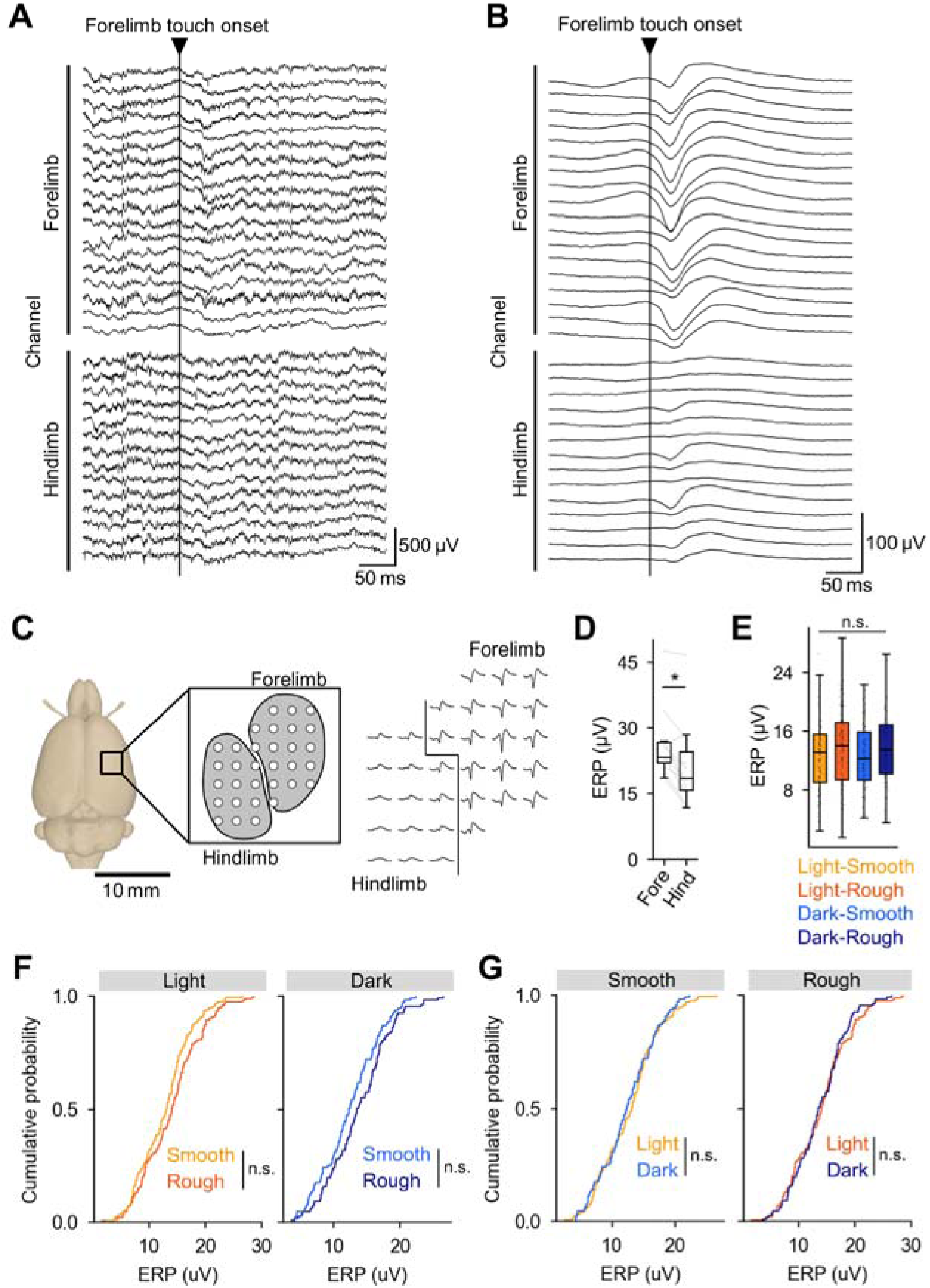
LFP recordings in rat S1 during walking. (**A**) A single representative LFP trace aligned to a forelimb contact. (**B**) An example of an averaged LFP trace from one session, aligned to forelimb contact. (**C**) The electrode montage and averaged LFP at each electrode. *Left*: The electrode montage showing all 32 recording sites. *Right*: Averaged LFP signals aligned to forelimb contacts with the floor, shown for each electrode depicted in the left panel. (**D**) Comparison of amplitudes between hindlimb and forelimb subregions, aggregated across all 11 rats. *P* = 1.89×10^-3^, *t*_10_ = -2.2, paired t-test, *n* = 11 rats. (**E**) Comparison of averaged amplitudes within the same trial for different floor textures and environmental conditions. *P* > 0.05, one-way analysis of variance (ANOVA) followed by TukeyLKramer post hoc test, *n* = 149, 149, 107 and 107 trials for smooth-light (light-orange), rough-light (dark-orange), smooth-dark (light-blue), and rough-dark (dark-blue), respectively. (**F**) Cumulative probability distributions of mean amplitude from each session, compared across different textures. *P* = 4.84×10^-1^, 8.35×10^-1^, *D* = 1.03×10^-1^ and 7.54×10^-2^ for light and dark environments respectively, two-sample KolmogorovLSmirnov test, *n* = 149 and 107 trials from 11 rats for light and dark, respectively. (**G**) Cumulative probability distributions of mean amplitude from each session, compared across light and dark environments. *P* = 1.05×10^-1^, 1.83×10^-1^, *D* = 0.14 and 0.149 for smooth and rough textures respectively, two-sample KolmogorovLSmirnov test, *n* = 149 and 107 trials from 11 rats for light and dark, respectively. Abbreviations: ERP, event-related potential; LFP, local field potential; S1, primary somatosensory cortex.

To see if the event-related responses were affected by the change in floor texture or the environmental condition, the amplitudes were compared. Despite the prominent negative deflection in LFPs at the forelimb contact with the floor, analyses showed no substantial differences in signal amplitude (Figure 3E, *P* > 0.05, one-way analysis of variance (ANOVA) followed by TukeyLKramer *post hoc* test, *n* = 149, 149, 107 and 107 trials for smooth-light, rough-light, smooth-dark, and rough-dark, respectively) when comparing rough *vs.* smooth textures (Figure 3F, *P* = 0.48 and 0.84, *D* = 1.03×10^-1^ and 7.54×10^-2^ for light and dark environments respectively, two-sample KolmogorovLSmirnov test, *n* = 149 and 107 trials from 11 rats for light and dark, respectively) and light *vs.* dark environments (Figure 6G, *P* = 0.11, 0.18, *D* = 0.14 and 0.149 for smooth and rough textures respectively, two-sample KolmogorovLSmirnov test, *n* = 149 and 107 trials from 11 rats for light and dark, respectively).

To further investigate, we computed grand-average LFP waveforms across combinations of floor texture (smooth *vs*. rough) and lighting condition (light *vs*. dark) (Figure 4A). Visual inspection of the mean waveforms revealed no distinct or prominent features. We then calculated channel-wise correlations between textures within each lighting condition (i.e., light-smooth *vs*. light-rough; dark-smooth *vs*. dark-rough). To assess whether lighting modulated texture-related correlations, we subtracted the correlation coefficients obtained in the dark from those in the light and averaged the differences across animals (Figure 4B). The correlation of LFP waveforms between different textures was stronger in the light condition, suggesting that in the dark environment, the average LFP waveform is more distinct. Additionally, we examined channel–channel correlation matrices across all 32 channels for each condition (Figure 4C). While stronger correlations were observed within individual S1 subregions, no systematic differences emerged between lighting conditions or between textures. To probe the frequency domain of the LFP, we computed time–frequency (wavelet) spectrograms across the floor texture and lighting condition combinations. However, no clear differences were observed across these combinations. Taken together, these results suggest that simple amplitude-based metrics may not fully capture subtler or higher-dimensional variations in underlying neuronal activity.

**Figure 4.**
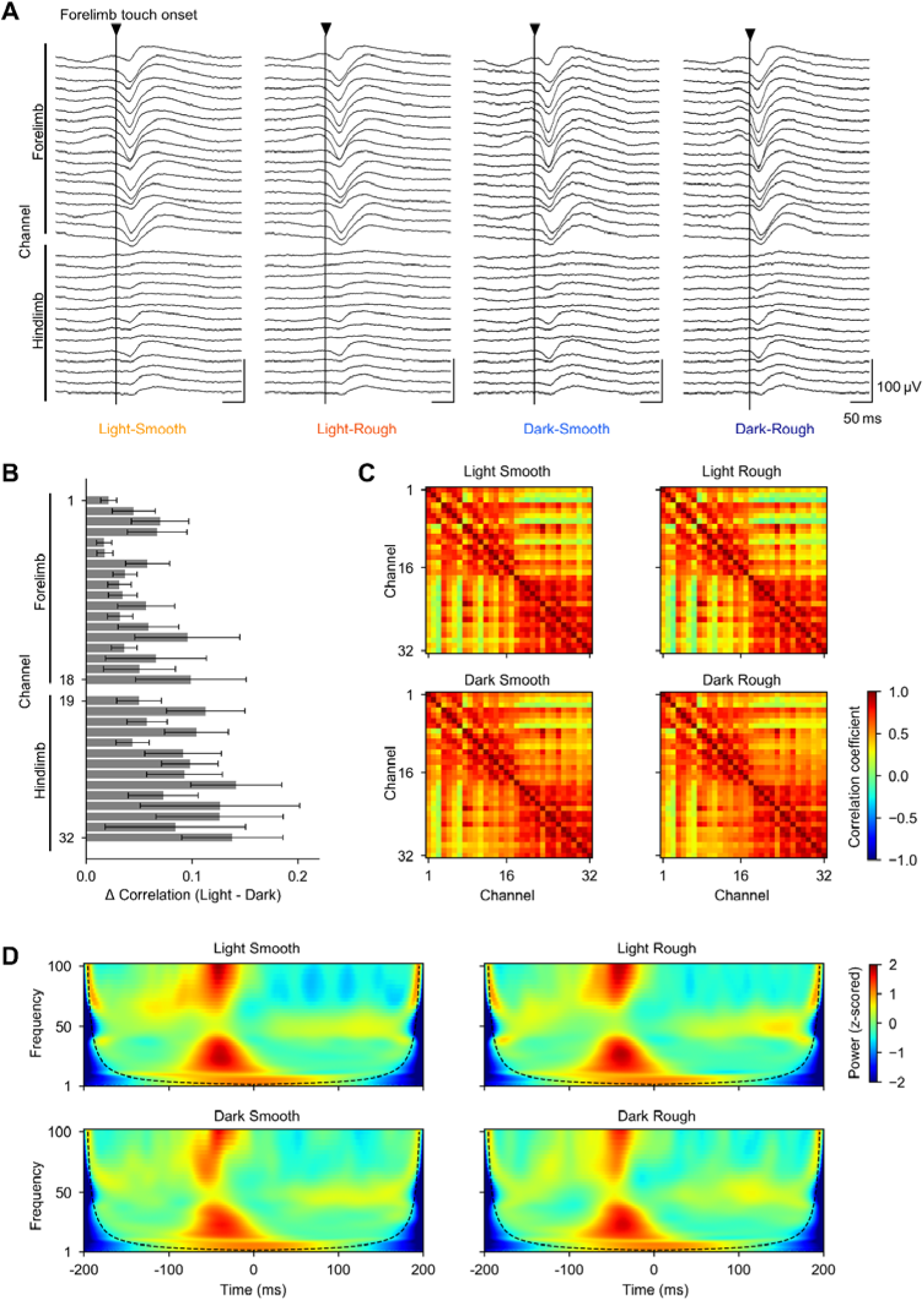
LFP characteristics across textures and lighting. (**A**) Example averaged LFP traces from one session, aligned to forelimb contact: light–smooth, light–rough, dark–smooth, dark–rough (left to right). (**B**) Channel correlation difference map (light − dark) computed from average LFP waveforms (Pearson’s *r*). Higher values indicate stronger channel-wise correlations in light relative to dark. (**C**) Correlation matrices shown separately for each texture and lighting condition. Matrices exhibit similar within–S1-subregion structure, with no clear texture- or light-dependent differences. (**D**) Example time–frequency (wavelet) spectrogram of LFP power from one session, aligned to forelimb contact. Dotted line indicates the cone of influence. Color scales represent wavelet coefficient magnitude (C) and corresponding power (D).

### Machine learning–based decoding of tactile and visual information

Given the absence of clear differences in averaged LFP features, we next examined how representations of textures in the LFPs varied across environments using machine learning approaches. We first implemented a support vector machine (SVM) classifier with a radial basis function (RBF) kernel. To reduce dimensionality, principal component analysis (PCA) was applied to the raw LFP traces (Supplementary Figure 2A). The SVM was then trained using 5-fold cross-validation, where, in each fold, 80% of the dataset was used to classify two types of labels: texture (smooth *vs*. rough) and trial (light *vs*. dark). Model performance was evaluated on the remaining 20% of the data. The overall classification accuracy was at chance level (∼50%), indicating that the PCA–SVM combination failed to extract informative features for texture or lighting condition from the LFP traces (Supplementary Figure 2B).

We next employed a deep learning model, a convolutional neural network (CNN), to classify both floor texture (smooth *vs*. rough) and trial (light *vs*. dark). The CNN architecture incorporated parallel convolutional pathways designed to capture both macro- and micro-scale temporal dynamics, followed by residual blocks. The network then branched into dual output layers to jointly predict texture and lighting conditions (Figure 5A).

**Figure 5.**
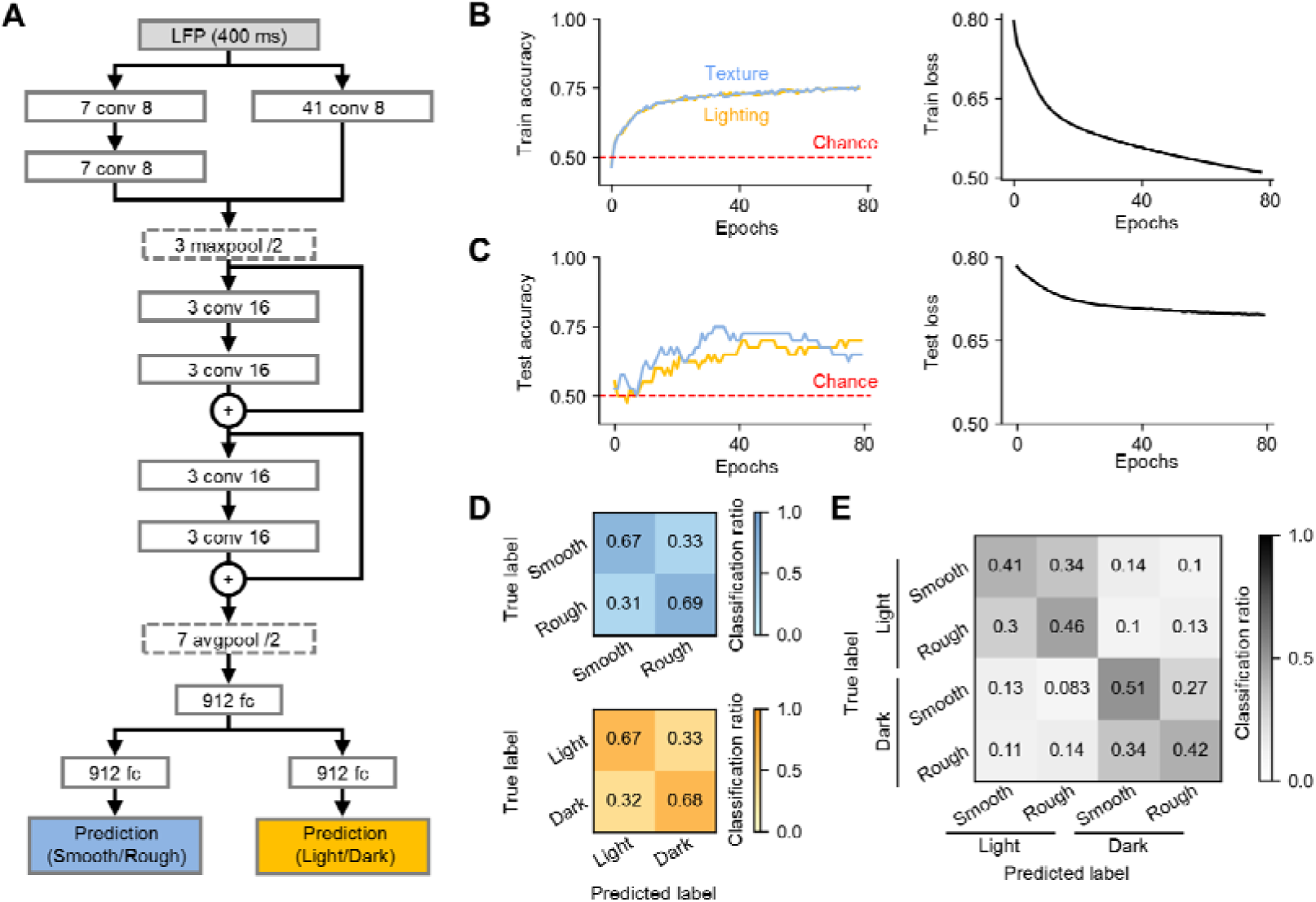
Model-based prediction of texture and environmental conditions from LFP. (**A**) The deep learning model architecture. The LFP input is processed through two parallel pathways for macro- and micro-scale feature extraction, followed by residual blocks that feed into two output heads for floor texture (Smooth *vs.* Rough) and the environmental condition (Light *vs.* Dark). (**B**) Training performance for a single representative rat. The left graph shows accuracy curves for texture (blue) and lighting (yellow), and the right graph shows the loss curves. (**C**) Testing performance for the same rat. The model exhibits good generalization, as accuracy increases and loss decreases on held-out data. (**D**) Confusion matrix for texture classification for all rats. Values above chance on the diagonal indicate successful texture prediction. Note that all values in the same row add up to 1. (**E**) Same as D, but for environmental conditions. (**F**) Combined confusion matrix for texture and trial predictions. The model performs well on both tasks across all rats. Abbreviations: avgpool, average pooling layer; conv, convolutional layer; maxpool, max pooling layer; LFP, local field potential.

Similarly to SVM, the model was trained and evaluated using 5-fold cross-validation. In training, the model exhibited stable learning curves (Figure 5B) and also generalized well to held-out test data (Figure 5C). Confusion matrices for both texture and lighting classifications were above chance along the diagonal, demonstrating that the model reliably extracted neural representations of the floor textures from the LFPs (Figure 5D–F).

### Neural representations become more distinct in darkness

To understand how the absence of the visual cue might refine tactile processing, we performed explainability analyses on the model’s learned representations (Figure 6). We extracted a 912-dimensional feature vector from the layer preceding the final outputs, then visualized these high-dimensional embeddings with scatter plots (Figure 6A, B). Clustering analyses using silhouette scores showed that representations of texture were more separated in the dark environment, suggesting that reduced visual input enhances the distinctness of neural dynamics that code tactile stimuli (Figure 6C, *P* = 2.31×10^-2^, *t*_8_ = -2.8, paired *t*-test, *n* = 9 rats, Supplementary Table 1). To account for trial order, silhouette scores were also calculated in sessions where dark trials preceded light trials. In this case, only an increasing trend was observed, with no significant difference detected (Supplementary Figure 3A, B *P* = 2.9×10^-1^, *t*_5_ = -1.16, paired *t*-test, *n* = 6 rats, Supplementary Table 2).

**Figure 6.**
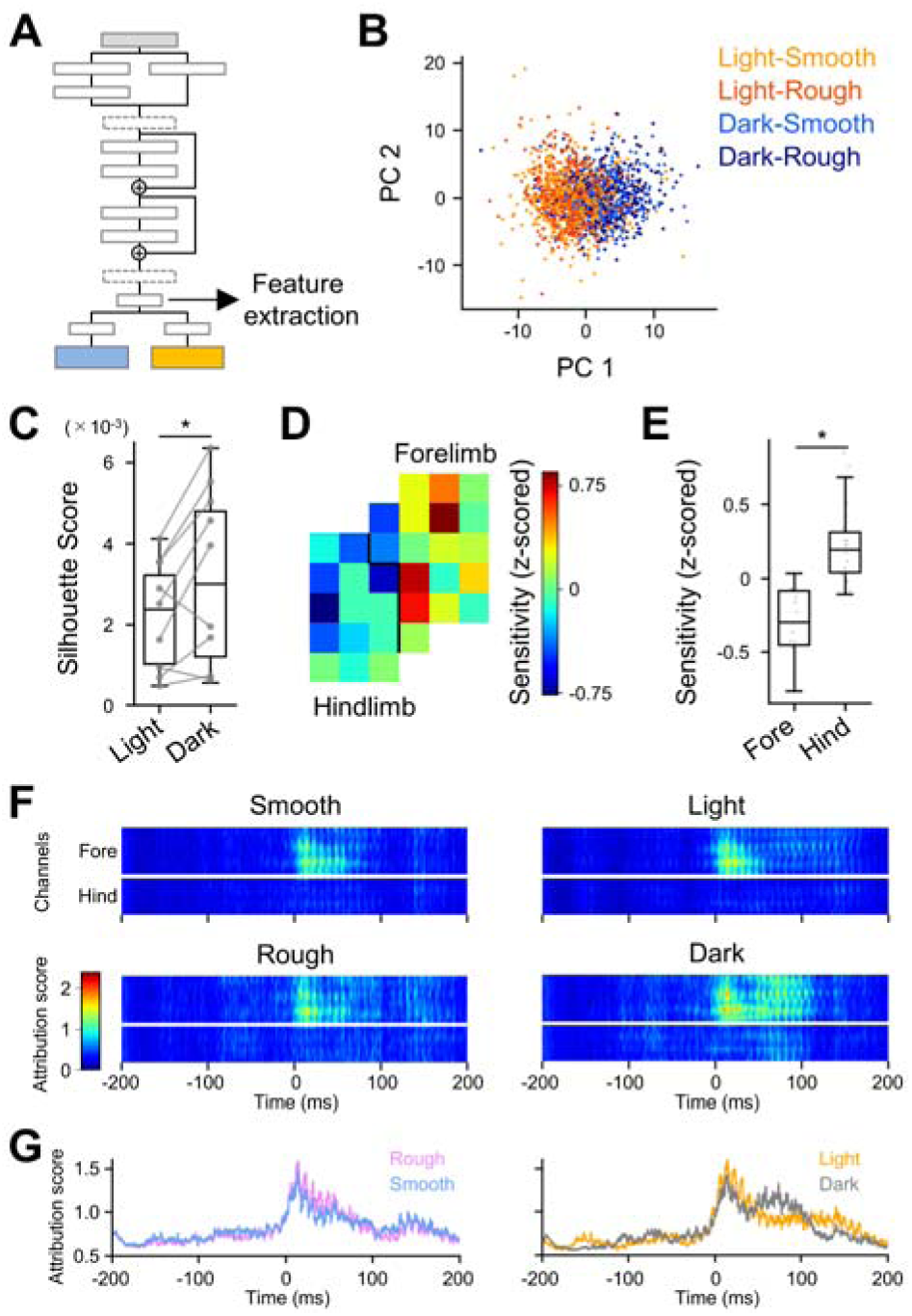
Neural representations are more distinct in dark environments than in light environments. (**A**) A 912-dimensional feature vector is extracted from the layer preceding the final output. (**B**) A scatter plot of these features from one rat shows individual LFP segments (aligned to forelimb contact). Light orange and dark orange correspond to the light conditions (smooth, rough), while light blue and dark blue correspond to the dark conditions (smooth, rough). (**C**) Silhouette scores across all nine rats, showing that the dark condition yields higher scores and thus more distinct neural representations. *P* = 2.31×10^-2^, *t*_8_ = -2.8, paired *t*-test, *n* = 9 rats. (**D**) A pseudo-color map based on occlusion analysis, illustrating the contribution of each electrode in the forelimb and hindlimb subregions. Hotter regions indicate higher importance for the model’s predictions. (**E**) Forelimb channels exhibit higher occlusion sensitivity than hindlimb channels, highlighting the forelimb’s dominant role when the foot contacts the floor. *P* = 4.53×10^-6^, *t*_16_ = -5.57, Student’s *t*-test, *n* = 9 rats. (**F**) Class activation maps generated via integrated gradients highlight key input features responsible for accurate model predictions of texture and environmental conditions. Attribution scores show each feature’s impact on the model’s output relative to a reference baseline: high positive scores denote features that strongly affect the predicted class. The onset of forelimb contact is aligned to time zero. (**G**) Attribution scores averaged over forelimb electrodes for floor texture (*left*) and the environmental conditions (*right*). A temporal lag in the dark condition suggests an extended processing window for floor texture when visual cues are absent.

To assess whether the observed differences in cluster separability between light and dark conditions could arise by chance, we performed a permutation test. Within each rat, the darkness/light labels were randomly shuffled across trials while preserving the number of trials per condition. For each permutation, silhouette scores were recalculated from the layer embeddings of the trained network. This procedure was repeated 1000 times to generate an empirical null distribution of cluster separability. The observed silhouette scores for the dark condition consistently exceeded the 95th percentile of the null distribution, confirming that the enhanced separability in darkness reflects systematic differences in neural representations rather than random variation (Supplementary Figure 4).

To elucidate which electrodes held the most information about the floor textures, we performed occlusion analysis. The results revealed that electrodes in the forelimb subregion contributed more to successful texture and lighting predictions (Figure 6D, E, *P* = 4.53×10^-6^, *t*_8_ = -5.57, Student’s *t*-test, *n* = 9 rats), which aligns with our observation that negative deflections in LFPs were largest in forelimb-targeting channels (Figure 2E). Further analysis using class activation maps from integrated gradients showed spatiotemporal patterns of salient features unique to each experimental condition (Figure 6F). In particular, the average attribution score in the forelimb subregion displayed a temporal shift in the dark condition, implying an extended processing window for texture information when visual cues are unavailable (Figure 6G, Supplementary Figure 5).

Collectively, these results indicate that visual deprivation modifies population-level activity in S1, yielding more distinctive representations of tactile stimuli. Such reorganization could provide a neural substrate for enhanced tactile perception under conditions of reduced or absent visual input.

## Discussion

This study provides evidence that S1 undergoes rapid reorganization of tactile representations when visual input is removed, even over a short period. Using high-density LFP recordings and deep learning techniques, we show that the neural encoding of tactile stimuli becomes more distinct under conditions of visual deprivation. Specifically, when rats were exposed to darkness, texture representations in S1 were more clearly distinguishable than under normal visual conditions. This demonstrates the adaptability of S1 and its capacity to rapidly adjust to changing sensory contexts.

A key finding was the significant increase in silhouette scores when texture representations were decoded in the dark. Silhouette scores, which quantify the separability of neural features, were notably higher under dark conditions, suggesting that visual deprivation sharpens the distinctions between tactile representations. While motivational factors, such as water restriction, could potentially confound the results, trial order reversals showed only a slight increase in silhouette scores during the dark condition. If motivation alone were responsible for the enhanced discriminability, silhouette scores should have been elevated during the later light trial in the reversed sequence. This supports the conclusion that the reorganization in S1 is more likely a direct result of visual deprivation rather than solely motivation or arousal. Although differences in silhouette scores were not significant in the reversed trials, this observation may reflect a lingering effect of prior visual deprivation, consistent with previous studies in humans showing that even brief periods of visual deprivation, such as blindfolding, can enhance tactile sensitivity. These findings suggest a robust and lasting effect of visual deprivation on tactile processing.

Our results also highlight the somatotopic specificity of tactile encoding in S1. The evoked potential in the forelimb subregion of S1 was significantly larger than in the hindlimb subregion, reflecting the topographic organization of S1, where distinct cortical areas process sensory inputs from different body parts (Ewert et al., 2008; Prsa et al., 2019; Sur et al., 1980). Further, occlusion analysis revealed that forelimb subregions were critical for distinguishing between tactile textures and lighting conditions. This finding underscores the dominance of the forelimb subregion in encoding tactile information, especially during the specific task of walking and contacting different textures. Integrated gradient analysis confirmed the importance of forelimb channels in encoding texture information and showed that the temporal window of processing following forelimb contact was extended in the dark compared to the light condition. Additionally, comparative analysis of the integrated gradient and time-frequency maps revealed a negative correlation in the 50-60 Hz frequency range, while a positive correlation was observed in the 80-90 Hz range. These findings suggest that frequency-specific patterns of LFP activity in different conditions are closely linked to the texture representations captured by the CNN model.

The extended temporal window in the dark condition suggests that visual deprivation may enhance the retention of tactile information in S1. Previous studies have shown that S1 neurons are involved in the short-term retention of tactile information (Zhou and Fuster, 2000, 1997, 1996), and prolonged neuronal firing in higher sensory areas may contribute to this sustained activity (Esmaeili and Diamond, 2019; Leavitt et al., 2017). Our findings suggest that the dark condition could lead to more prolonged neural representations of tactile stimuli. Whether this reflects a compensatory mechanism for the absence of visual input or a correlation with heightened tactile sensitivity requires further exploration.

Environmental illumination can also influence the arousal state, which in turn affects cortical dynamics as reflected in LFPs. Dark environments typically promote exploratory behavior and increase arousal, modulating cortical activity through irradiance- and cone-opponent–dependent pathways to arousal circuits like the locus coeruleus and basal forebrain (Tamayo et al., 2023). Arousal in S1 enhances both baseline activity and stimulus encoding, with neuromodulatory drive sharpening temporal precision (Eggermann et al., 2014; Poulet and Petersen, 2008; Shimaoka et al., 2018). These state-dependent effects may account for the enhanced separability of tactile representations observed in the dark, despite minimal differences in average evoked amplitude (Lee et al., 2020; McGinley et al., 2015; Shimaoka et al., 2018; Vinck et al., 2015). The shift toward a desynchronized, high-arousal state in darkness likely reduces low-frequency shared variability and boosts fast-timescale signal components, improving the separability of neural clusters in a high-dimensional LFP space. Moreover, neuromodulatory engagement during heightened arousal can extend effective integration windows and enhance gain in task-relevant networks, consistent with the longer temporal window for accurate predictions seen in forelimb channels.

These findings highlight the limitations of traditional analysis methods, such as amplitude-based metrics and event-related potentials, which often fail to capture subtle, higher-dimensional features in neural signals (Saxena and Cunningham, 2019; D. L. Yamins and DiCarlo, 2016; D. L. K. Yamins and DiCarlo, 2016). In contrast, our deep-learning approach revealed fine-grained spatiotemporal patterns in LFPs, showcasing enhanced texture-specific separability under visual deprivation. This demonstrates the power of advanced computational tools to uncover previously inaccessible shifts in sensory coding, with broad applicability to other high-dimensional neural datasets, such as those from multi-electrode arrays in freely behaving animals. However, it’s important to note that each model was trained specifically for each rat, and inter-animal generalization remains a challenge, due to differences in electrode placement and individual brain structure. Despite this limitation, the model’s ability to extract complex, high-dimensional patterns from the dataset remains evident, and the results show significant progress in detecting sensory changes.

While fMRI studies have demonstrated tactile enhancement and corresponding neural reorganization during short-term visual deprivation (Facchini and Aglioti, 2003; Kauffman et al., 2002; Pascual-Leone and Hamilton, 2001b), the low temporal and spatial resolution of fMRI limits its ability to capture detailed changes in neural representations. Our study, through LFP recordings, provides a more nuanced picture of the population-level activity underlying tactile perception enhancement in the absence of visual input. Future studies could extend this work to assess whether rats can discriminate textures more accurately in the dark and explore the direct relationship between neural coding and perceptual performance.

From an evolutionary perspective, enhanced tactile sensitivity in low-visibility environments provides an adaptive advantage. Many species, including rodents, rely on somatosensation for navigation and foraging when visual information is scarce. The ability to rapidly enhance tactile processing in such conditions could aid in efficient resource acquisition and predator detection. Given the influence of arousal on tactile sensitivity (Lee et al., 2020; Shimaoka et al., 2018), the dynamic nature of sensory processing becomes evident—S1 does not merely respond passively to tactile input but actively adapts its encoding strategies based on available sensory cues.

Several limitations warrant further exploration. First, if arousal contributes to the effects observed in S1 during light–dark transitions, we predict that arousal indices, such as pupil or whisking activity, should correlate with LFP spectral composition and decoding accuracy on a trial-by-trial basis (McGinley et al., 2015; Reimer et al., 2014; Shimaoka et al., 2018). Second, manipulating neuromodulatory tone through optogenetic activation of arousal circuits should modulate the separability of texture representations, independent of illumination (Eggermann et al., 2014; Harris and Thiele, 2011). Future experiments integrating real-time arousal measures and LFP analyses will be crucial in disentangling the contributions of arousal versus light–dark effects on cross-modal reorganization in S1. Additionally, while rats serve as a powerful model for sensory processing, their neural architecture may not fully capture the complexities of human perception. Extending this paradigm to assess perceptual changes in tactile acuity will further clarify the relationship between neural coding and perceptual performance.

In conclusion, our study provides new insights into the brain’s remarkable ability to reorganize its sensory processing in response to changes in sensory input. By demonstrating that visual deprivation rapidly reconfigures tactile processing in S1, we highlight the brain’s flexibility in adapting to environmental changes. This work underscores the potential of combining behavioral paradigms, LFP recordings, and deep learning techniques to understand the dynamic and adaptive nature of sensory coding, paving the way for future research in multisensory interactions.

## Data availability

The data that support the findings of this study are available from the corresponding author upon request.

## Code availability

Custom code generated during this study for data analysis are available at https://github.com/UT-yakusaku/Yamashiro-eLife-2024.

## Acknowledgments

This work was supported by JST ERATO (JPMJER1801), AMED-CREST (24wm0625401h0001; 24wm0625502s0501; 24wm0625207s0101; 24gm1510002s0104), the Institute for AI and Beyond of the University of Tokyo, JSPS Grants-in-Aid for Scientific Research (18H05525, 20K15926, 22K21353, 22J22097), KOSÉ Cosmetology Research Foundation, the Public Foundation of Chubu Science and Technology Center, and Konica Minolta Science and Technology Foundation.

