## Supplementally Materials for "Enhanced Tactile Coding in Rat Neocortex Under Darkness"

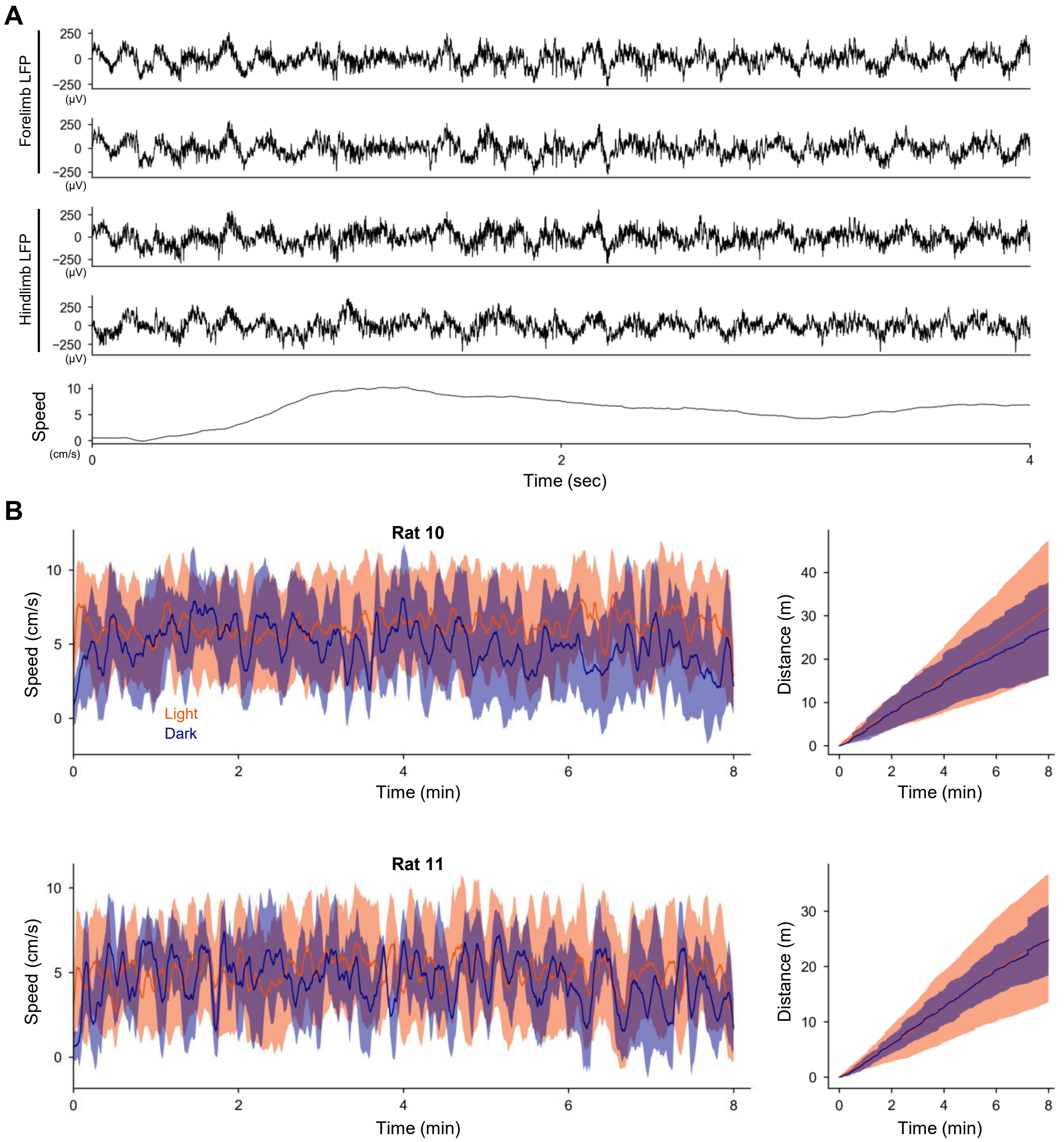


**Supplementary Figure 1. Locomotor speed traces during each trial**

(**A**) Locomotor-speed traces aligned with corresponding LFPs. (**B**) Locomotor speed traces across trials for two rats. *Top left*: Locomotor speed trace for Rat 10, with the solid line representing the mean and the shaded area indicating the standard deviation (Orange: light-trial, Navy: dark-trial). *Top right*: Cumulative distance traveled during trials for Rat 10, with the solid line representing the mean and the shaded area indicating the standard deviation (Orange: light-trial, Navy: dark-trial). *Bottom left and right*: Same as the top row but for Rat 11.

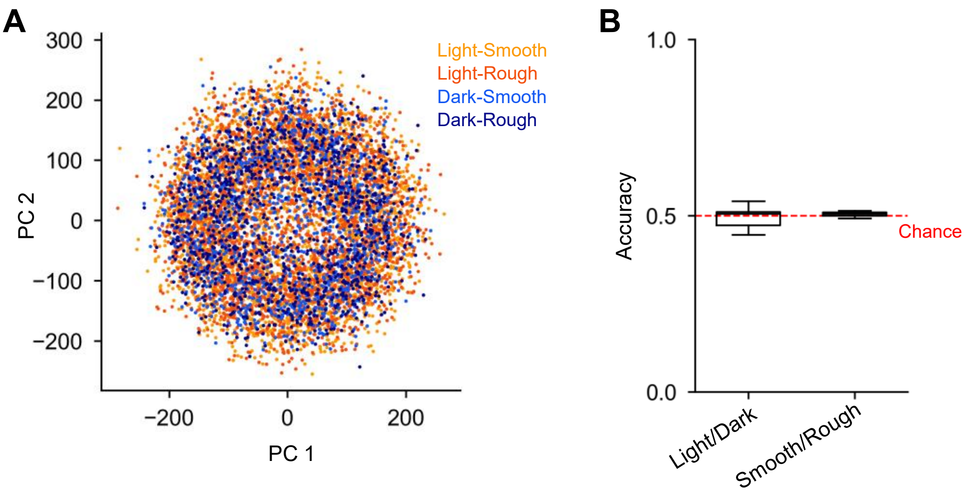


**Supplementary Figure 2. LFP classification using PCA and SVM**

(**A**) LFP traces aligned to forelimb onset, mapped to two-dimensional space using PCA. Light orange and dark orange represent the light conditions (smooth, rough), while light blue and dark blue represent the dark conditions (smooth, rough). (**B**) Classification accuracy for two categories (light *vs*. dark and smooth *vs*. rough) using an SVM classifier. The red dotted line indicates the chance level.

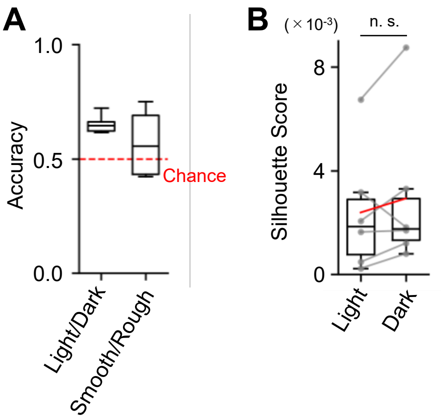


**Supplementary Figure 3. Separability of texture representations in inverted session**

(**A**) Silhouette scores across six rats during sessions where the dark trial preceded the light trial.
(**B**) Classification accuracy of LFPs recorded during sessions where the dark trial preceded the light trial.

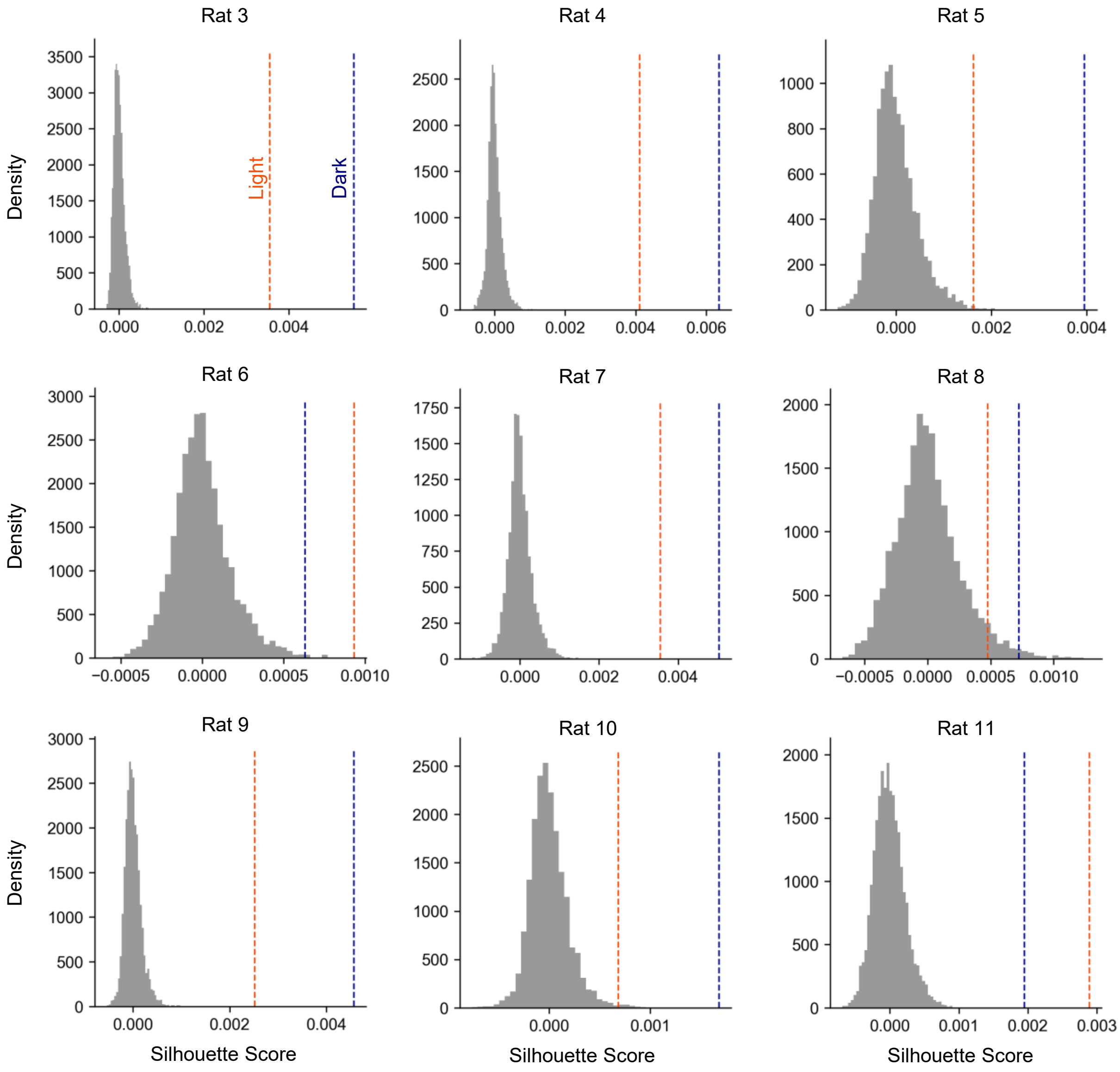


**Supplementary Figure 4. Silhouette score for each individual rat**

Each panel shows the empirical null distribution and silhouette scores for light and dark trials for each rat. The gray histogram represents the null distribution, with the orange dotted line indicating the silhouette score for light trials and the navy dotted line representing the silhouette score for dark trials.

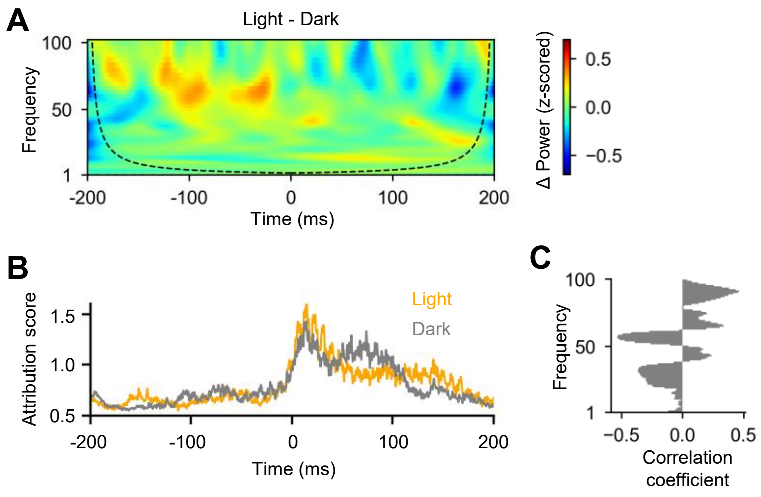


**Supplementary Figure 5. Spectral analysis of leaned features by CNN model**

(**A**) Pseudo-colormap showing the differences in averaged wavelet transform results between light and dark conditions of aligned LFPs at forelimb onset. Dotted line indicates the cone of influence. (**B**) As in Figure 6G (right panel), attribution scores averaged over forelimb electrodes for the environmental conditions. (**C**) Correlation coefficient between the difference in attribution scores and the difference in frequency elements within the spectral map, calculated for each frequency element.

| Rat # | Silhouette Score (mean ± SD) | |
|---|---|---|
| | **Light (mean ± SD)** | **Dark (mean ± SD)** |
| Rat 1 | - | - |
| Rat 2 | - | - |
| Rat 3 | $3.55\times10^{-3} \pm 8.25\times10^{-4}$ | **$5.52\times10^{-3} \pm 2.08\times10^{-3}$** |
| Rat 4 | $4.12\times10^{-3} \pm 3.70\times10^{-4}$ | **$6.36\times10^{-3} \pm 9.07\times10^{-4}$** |
| Rat 5 | $1.62\times10^{-3} \pm 5.74\times10^{-4}$ | **$3.95\times10^{-3} \pm 2.35\times10^{-3}$** |
| Rat 6 | **$9.33\times10^{-4} \pm 3.20\times10^{-4}$** | $6.32\times10^{-4} \pm 4.67\times10^{-4}$ |
| Rat 7 | $3.55\times10^{-3} \pm 5.43\times10^{-4}$ | **$5.03\times10^{-3} \pm 9.93\times10^{-4}$** |
| Rat 8 | $4.77\times10^{-4} \pm 4.32\times10^{-4}$ | **$7.23\times10^{-4} \pm 5.29\times10^{-4}$** |
| Rat 9 | $2.52\times10^{-3} \pm 4.63\times10^{-4}$ | **$4.57\times10^{-3} \pm 1.25\times10^{-3}$** |
| Rat 10 | $6.83\times10^{-4} \pm 5.18\times10^{-4}$ | **$1.68\times10^{-3} \pm 4.87\times10^{-4}$** |
| Rat 11 | **$2.89\times10^{-3} \pm 3.14\times10^{-4}$** | $1.95\times10^{-3} \pm 3.30\times10^{-4}$ |

**Supplementary Table 1. Silhouette score in light and dark trials for each rat**
Larger values are indicated in bold.

| Rat # | Silhouette Score (mean ± SD) | |
|---|---|---|
| | Light (mean ± SD) | Dark (mean ± SD) |
| Rat 1 | **$3.17\times10^{-3} \pm 1.08\times10^{-3}$** | $1.82\times10^{-3} \pm 7.19\times10^{-4}$ |
| Rat 2 | $2.34\times10^{-4} \pm 1.27\times10^{-3}$ | **$8.01\times10^{-4} \pm 8.99\times10^{-4}$** |
| Rat 3 | $1.64\times10^{-3} \pm 1.23\times10^{-3}$ | **$1.69\times10^{-3} \pm 6.16\times10^{-4}$** |
| Rat 4 | $2.06\times10^{-3} \pm 5.84\times10^{-4}$ | **$3.30\times10^{-3} \pm 5.78\times10^{-4}$** |
| Rat 5 | $6.74\times10^{-3} \pm 2.59\times10^{-3}$ | **$8.74\times10^{-3} \pm 4.32\times10^{-3}$** |
| Rat 6 | $4.79\times10^{-4} \pm 1.17\times10^{-3}$ | **$1.21\times10^{-3} \pm 1.03\times10^{-3}$** |

**Supplementary Table 2. Silhouette score in light and dark trials for each rat in inverted session**

Larger values are indicated in bold.
